# Hierarchical Gaussian Processes and Mixtures of Experts to Model COVID-19 Patient Trajectories

**DOI:** 10.1101/2021.10.01.462821

**Authors:** Sunny Cui, Elizabeth C. Yoo, Didong Li, Krzysztof Laudanski, Barbara E. Engelhardt

## Abstract

Gaussian processes (GPs) are a versatile nonparametric model for nonlinear regression and have been widely used to study spatiotemporal phenomena. However, standard GPs offer limited interpretability and generalizability for datasets with naturally occurring hierarchies. With large-scale, rapidly-updating electronic health record (EHR) data, we want to study patient trajectories across diverse patient cohorts while preserving patient subgroup structure. In this work, we partition our cohort of over 2000 COVID-19 patients by sex and ethnicity. We develop and apply a hierarchical Gaussian process and a mixture of experts (MOE) hierarchical GP model to fit patient trajectories on clinical markers of disease progression. A case study for albumin, an effective predictor of COVID-19 patient outcomes, highlights the predictive performance of these models. These hierarchical spatiotemporal models of EHR data bring us a step closer toward our goal of building flexible approaches to capture patient data that can be used in real-time systems^*^.

## 1. Introduction

The highly contagious nature of the emergent coronavirus (COVID-19) and limited knowledge of treatment methods necessitate decision support tools that can efficiently estimate and predict patient trajectories in order to measure disease progression. Notably, recent findings report considerable disparities in manifestations of COVID-19 across racial minorities within the United States, with a disproportionately high frequency of hospitalizations among African American, Hispanic, and Native American populations.^1^ Higher rates of obesity, a known high-risk comorbidity, are observed in marginalized groups, which contribute to more severe illnesses and higher mortality rates for these patients.^2^ Worse outcomes arise due to a complex combination of physiological, socioeconomic, behavioral, and cultural factors. A model that can account for group structures that arise both inherently and environmentally is necessary in order to develop clinical recommendations tailored to individual patients and to mitigate bias in treatment procedures; at the same time, that model should also allow for the sharing of signal across groups when patient group sample sizes are small.

The Hospitals at the University of Pennsylvania (HUP) COVID-19 dataset contains clinical observations of 2069 patients who tested positive for COVID-19 via a PCR test between April 2020 to August 2020 at the University of Pennsylvania Medical Center (UPMC) hospital in Philadelphia, PA.

This anonymized dataset includes the following patient information:

- patient demographic information including age, sex and ethnicity;
- labs and vital sign measurements, including blood serum creatinine, partial pressure of oxygen, and total urine output;
- procedural information, including details of mechanical ventilation, nasal cannula, and liters of oxygen flow; and
- medication information including type, dosage, and time of administration.

With an emergent disease like COVID-19, we want a model that is robust to missing and noisy patient data, and also computationally tractable to allow continuous data updates. Known for their flexibility, interpretability, and uncertainty quantification, Gaussian processes (GPs) have proven useful in machine learning,^3^ spatiotemporal statistics,^4^ and functional data analysis.^5^ Among their applications, GP regression is a nonparametric regression model that places a distribution on arbitrary nonlinear functions with smoothness modulated by the selected kernel function.^6^ Updated by observations, the GP posterior enables predictions and uncertainty estimates at unobserved locations on sequences, such as the time or space domain, including the future. Due to the Gaussian assumption of the joint distributions over observations, the posterior is Gaussian with closed-form mean and variance terms.

Previous work has exploited the flexibility of GPs to obtain insights into problems in healthcare, including early detection of sepsis through multi-output GPs,^7,8^ online updates of patient vital signals with sparse multi-output GPs,^9^ and reliable prediction of adverse hospital events by jointly modeling longitudinal trajectories and time-to-event data.^10^

For the task of modeling disease trajectories, particularly for a large patient cohort, using standard GP regression is insufficient because many complex diseases such as lupus and pneumonia manifest heterogeneously in patients across different demographic and clinical sub-groups.^11,12^ Noting this heterogeneity, prior work placed a hierarchy on scleroderma patients at the population, subgroup, and individual levels.^10^ B-splines were used to model each subgroup trajectory and a GP was used to capture noise.

Although the MedGP approach^9^ combined information across patients using an empirical Bayes approach, allowing subgroups to be captured via kernel parameters, it lacks a rigorous approach to evaluating group structure and posteriors. Motivated by the need to explicitly account for group structure, our framework builds on the premise of a group structure in the patient population and provides a fully Bayesian treatment of hierarchical disease trajectory modeling.

The contributions of this work are as follows: At a high level, we develop a flexible Gaussian process that is able to capture sparse, noisy, electronic health record (EHR) time-series data. More specifically, we build a hierarchical mixture of experts (MOE) Gaussian process (GP) regression model that allows sharing of strength across patient samples with known group structure. The MOE allows each sample to participate in multiple patient groups simultaneously, such as inclusion in both the female (sex) and Black (race) patient groups. Furthermore, our fast closed-form inference method allows us to apply this framework to hundreds of COVID-19 patient trajectories to show its robustness in fitting a variety of clinically important covariates.

This paper is organized as follows: In Section 2, we discuss the background for standard, hierarchical, and MOE Gaussian process regression models. We introduce our framework of MOE hierarchical Gaussian process regression in Section 3. We demonstrate the performance of our framework on COVID-19 patient EHR data and discuss the implications of these results in Section 4. We conclude by exploring future directions in Section 5.

## 2. Background

In this section, we provide a brief summary of GP regression and its extension to a Bayesian hierarchical setting.

### 2.1. Gaussian process regression (GPR)

We consider the Bayesian analysis of standard linear regression *f* (*x*_*i*_) = *β*^*T*^ *x*_*i*_, where *β* is the weights of the linear model, *x*_*i*_ are regressors, and *f* (*x*_*i*_) is the noiseless function. Given observed data 𝒟 = (*X, Y*) where 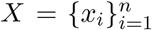 are regressors such as time across *n* total observations, and 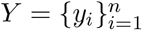 are noisy, scalar responses, then we can write each response as *y*_*i*_ = *f* (*x*_*i*_)+ ϵ_*i*_ where ϵ_*i*_ ∼ 𝒩 (0, *σ*^2^) is Gaussian white noise. Given a new set of regressors *X*_∗_ = {*x*_∗_}, the goal is to predict the responses *Y*_∗_ = *f* (*X*_∗_).

We can extend these linear models to nonlinear regression functions using Gaussian processes. Gaussian process regression is a probability distribution over arbitrary smooth functions such that any finite realization is a multivariate Gaussian random variable. For any observations *X* = [*x*_1_, ···, *x*_*n*_],

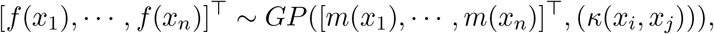

where *m*(·) is the mean function and *κ*(·, ·) is a positive definite kernel function. As in prior work, the mean function *m* is assumed to be zero.^9^ There are many possible positive definite kernel functions *κ*, including exponential (Ornstein-Uhlenbeck), squared exponential, and Matérn covariance functions. These covariance functions include parameters that control the spatial variance and decay of the dependency over the domain; these kernel parameters are often estimated by maximizing the log likelihood (MLE):

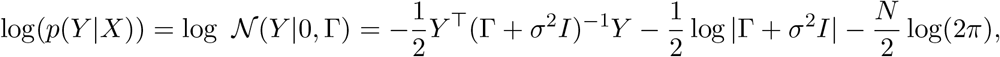

where Γ_*ij*_ = *κ*(*x*_*i*_, *x*_*j*_). Let Γ_∗_ = *κ*(*X, X*_∗_) and Γ_∗∗_ = *κ*(*X*_∗_, *X*_∗_) then the posterior of *Y*_∗_ is given by

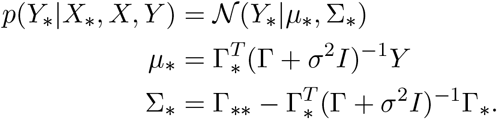

A point estimate of *Y*_∗_ is given by *μ*_∗_, the posterior mean, while Σ_∗_ is the variance of this posterior mean.

The computational complexity of inference for GPR is O(*n*^3^) because of the need to invert Γ, an *n* by *n* matrix. Fortunately, there is an immense literature on scalable inference algorithms for GPs, including tapering.^13^ The idea of tapering is to impose zero correlation between two points that are not close to each other by multiplying *κ* by a tapering function *T* : *κ*_*T*_ := *κ*(*x, y*)*T* (*x, y*). For example, when *T* (*x, y*) = 1_*{*‖*x*−*y*‖*<ϵ}*_, *κ*_*T*_ (*x, y*) = 0 if ‖*x* − *y*‖ ≥ ϵ, resulting in a sparse block diagonal covariance matrix.

### 2.2. Hierarchical Gaussian process (HGP) regression

One of the main challenges in predicting future values of a disease trajectory or imputing unobserved values within a trajectory is that biological and environmental factors lead to high variance in patient state and disease progression. For instance, many diseases include one or more disease subtypes, and the progression and severity of a disease can vary across patients with different ages, sexes, or chronic conditions.

For datasets with known subgroups, hierarchical models are a natural choice because they allow the sharing of information across and within subgroups. The use of hierarchical models allows precise modeling of each subgroup and sharing of signal across all of the subgroups; it is particularly beneficial in the case where each subgroup has a small sample size.

Hierarchical structure can be enforced through the mean function, the covariance function, or a structured prior. Prior work [14] placed a hierarchy on the mean function parameters to model *PM*_2.5_ levels, a measurement of air quality, much like the spline model for individualized disease prediction.^10^ Other work [15] placed a hierarchy on gene expression at two levels— each experiment and each replicate gene—to model heterogeneity. Conjugate inverse Gamma priors were placed on the kernel parameters to model the relationships between low and high accuracy experiments.^16^ Variants of the hierarchical model include hierarchical MOE that lends a tree structure in computing parameter values,^17^ deep GPs in which inputs to each GP have their own GP prior.^18^ This work uses subsets of inducing points to fit experts, which hold information at the group and individual levels.^19^

## 3. Hierarchical Gaussian process regression for patient trajectories

In the context of prior work, we develop a Bayesian hierarchical GP regression model for patient data. We group the patient population by attributes including sex and ethnicity. We impose a hierarchy on these trajectories at the group and individual levels by letting the mean of each level in the hierarchy be distributed by a Gaussian process parameterized for the level above. We use *k* = 1, ···, *K* as the group-level subscript and *i* = 1, ···, *N*_*k*_ as the patientlevel subscript in group *k*. All patients in the *k*th subgroup share an underlying trajectory modeled by *g*_*k*_(*x*). Patient *i* in subgroup *k* is associated with a unique trajectory, denoted by *f*_*k,i*_(*x*), that is influenced by various factors including demographics, lifestyle choices, genetic predispositions, and pre-existing conditions. Then,

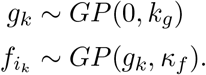

Let 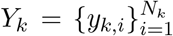 be the collection of noisy observations of clinical markers of *N*_*k*_ patients in subgroup *k* at time points 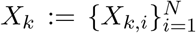. The covariance between the data *Y* and the functions *f* (·), *g*(·) is

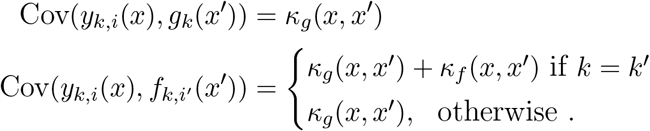

### 3.1. HGP kernel functions and tapering

Our model uses an additive hierarchical kernel, similar to that introduced by [15], with tapering that further enforces sparsity. For flexibility in the smoothness of the inferred functions, we choose the Matérn kernel with parameter *ν* that controls the smoothness of the GP:^20^

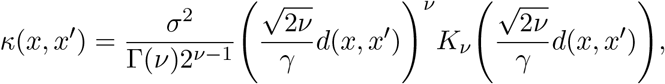

where *K*_*ν*_ is the modified Bessel function of the second kind with order *ν*. In practice, we estimate these parameters by maximizing the likelihood. In our model, we set kernel parameter 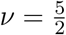 at the group level, and 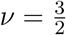 at the individual level.

With this kernel function, we model the data distribution as multivariate normal.

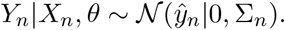

The parameters *θ* are {*α*^*T*^, *β*^*T*^, *γ*^*T*^} . The covariance matrix Σ_*n*_ is written as

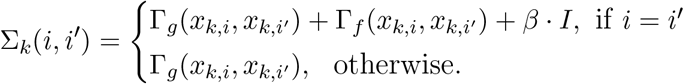

Both Γ_*g*_ and Γ_*f*_ are matrices formed by evaluating *κ*_*g*_ and *κ*_*f*_, respectively, on *x*_*k,i*_ and *x*_*k,i′*_. These covariance matrices inherit a natural block structure from the kernels (Fig. 2). To scale up the HGP with computational complexity O(*n*^3^), we further perform tapering to enforce relationships only between close time points. Tapering encodes sparsity in the covariance matrix on the off-diagonal elements that are more distant from each other in time, which improves inference tractability.^13^

**Fig. 1:**
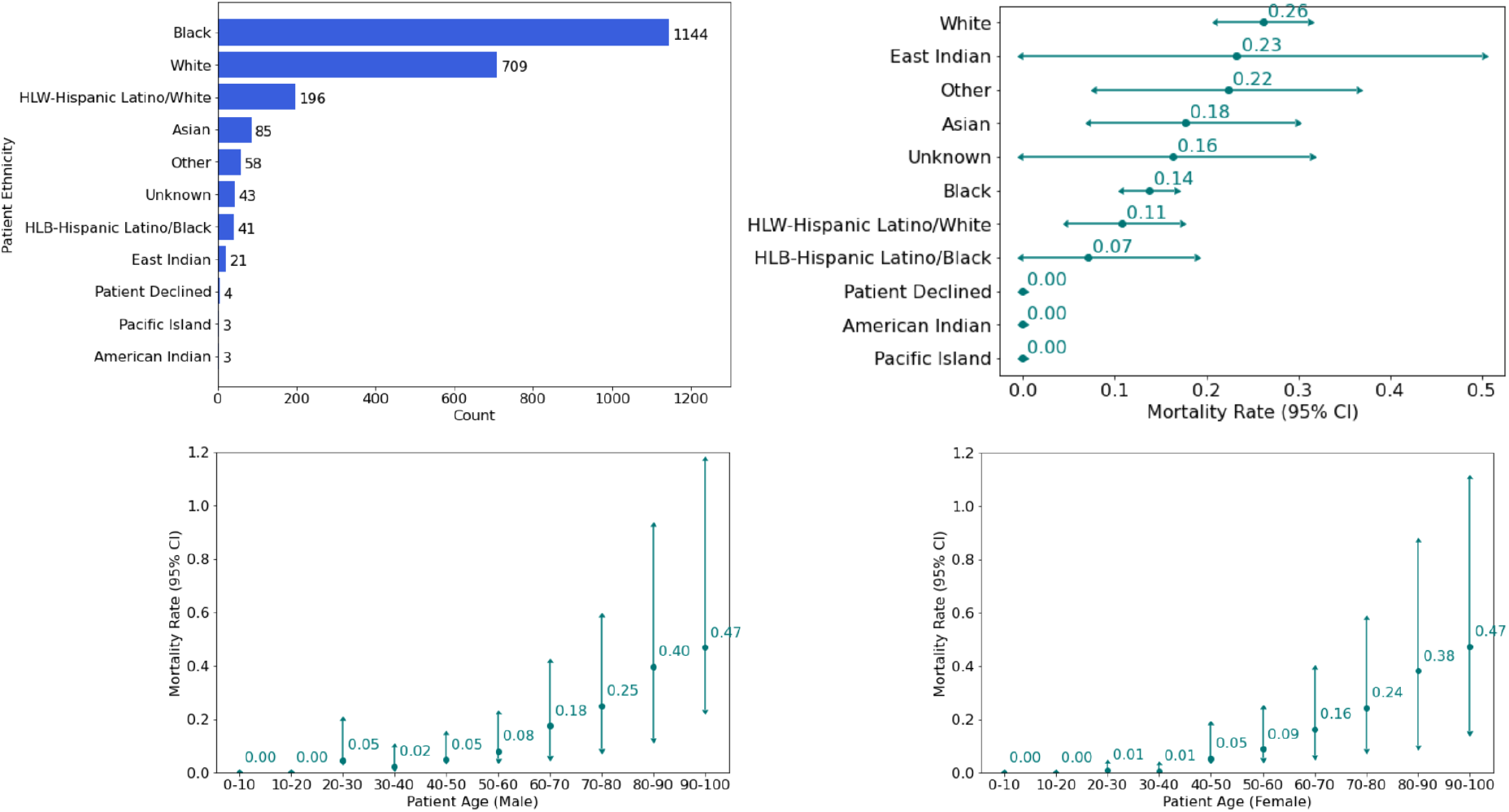
Patient cohort breakdown. Cohort size (top left); patient mortality by ethnicity (top right); patient mortality by age and sex (males bottom left and females bottom right)

**Fig. 2:**
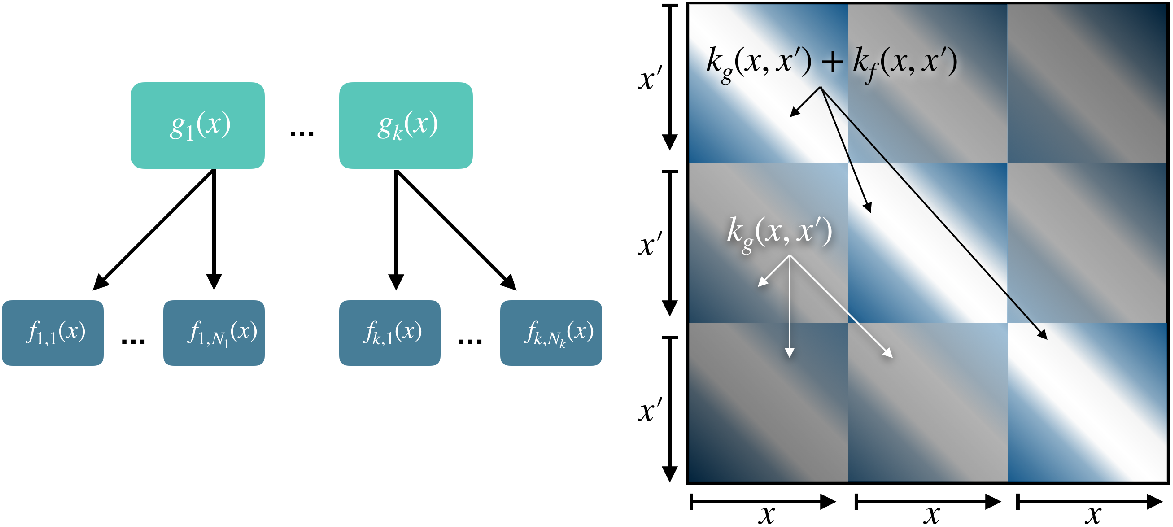
Model setup (left) and block structure of HGP covariance matrix (right).

### 3.2. Mixture of experts

Although the HGP allows us to model group structure and individual patient trajectories that differ from the group, its exponential cost with respect to number of groups renders it impractical for large patient cohorts with many groups. Because each patient belongs to multiple groups simultaneously – sex, ethnicity, and disease subtype for instance – we want a tractable way to combine information from all of the patient’s group attributes, i.e., an additive kernel. Thus, we extend the HGP with mixture of experts (MOE) kernels at the group level (Fig. 3). Originally developed to handle multiple modalities in large datasets,^21^ MOE GPs can be adapted to a hierarchical setting such that the group-level kernel is the sum of attribute kernels of patients belonging to that group. An ensemble of local *experts* allows the kernel function to adapt to each observation,^22^ which in our case corresponds to a patient. Again, we use a tapered Matérn 5/2 kernel at the group level and a tapered Matérn 3/2 kernel at the patient level. We perform efficient close-form inference using the SciPy Optimizer.

**Fig. 3:**
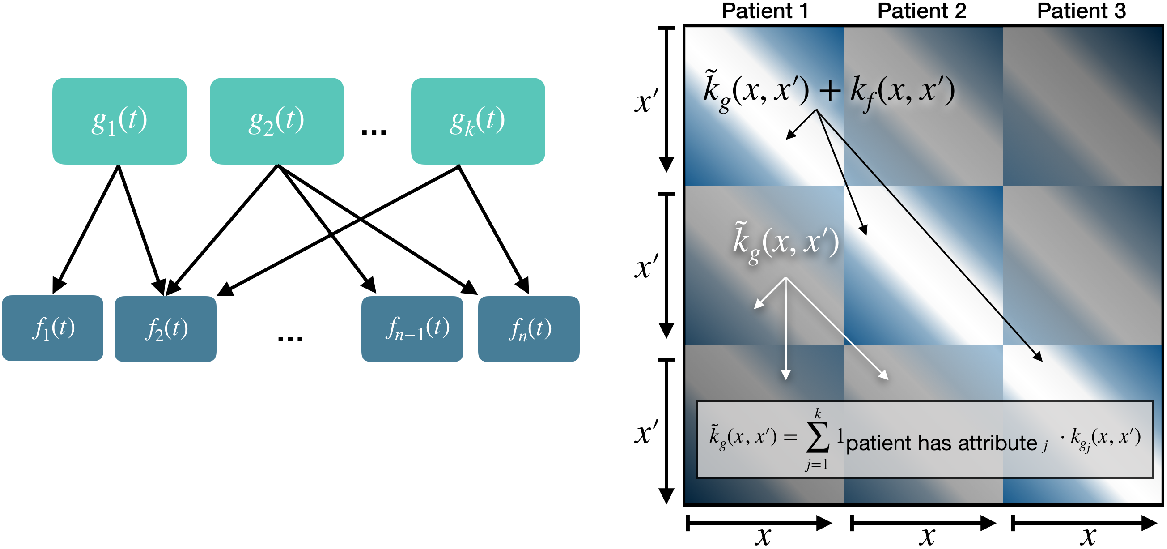
Model setup (left) and MOE HGP covariance matrix (right).

## 4. Experiments

We first benchmark our MOE HGP model, using HUP patient trajectories, against standard GPR and an HGP. We then present examples of fitted and predicted trajectories of cluster representatives, or patients whose trajectory minimizes the Wasserstein distance to all other patients in their subgroup. Intuitively, the cluster representative corresponds to the patient who best captures the canonical trajectory of that group.

We evaluate the performance of our MOE HGP on COVID-19 patient trajectories from the Hospitals at the University of Pennsylvania (HUP). For the purposes of model fitting, we only consider patient trajectories with over 25 observations corresponding to unique time points. We group patients based on attributes of sex (*male* and *female*) and ethnicity (*Black* and *white*). We create balanced patient cohorts with 30 patients per permutation of groups (i.e., 30 Black women, 30 Black men, 30 white women, 30 white men).

For each patient and each covariate, we select 25% of the measurements randomly as the test set and use the remaining measurements as the training set. It is also possible to include future time points in the test set, albeit at the expense of GP model performance as test points extend further into the future, meaning there is greater uncertainty in the predictions.^20^ To evaluate performance, we use mean squared error (MSE) and *R*^2^ metrics to compare the train and test sets to predicted values.

We also evaluate the 95% confidence intervals (CIs) to measure model calibration for GPR, HGP, and MOE HGP. In our discussion, the values reported for 95% CI calibration refer to the percentage of points that fall outside the 95% confidence interval. We focus on *albumin* as our covariate of interest, as it has been shown to be a clinical marker of COVID-19 progression.^23^ The results for *albumin* are representative of trends across covariates in the dataset (see Supplementary material for details).

The shapes of the patient trajectories for albumin vary greatly (Fig. 4). GPR cannot, for example, capture the trajectory of patient 11, but the HGP and MOE HGP are able to do so. For patient 38, the more granular trends for the first few time points are captured by the MOE HGP, but not the HGP. The average train MSE across patients for the covariate albumin is the lowest for the MOE HGP. The average test MSE across patients is comparable across the three models. However, the train and test *R*^2^ values, and the 95% CI calibration, are much better across patients for the HGP and MOE HGP as compared to GPR (Table 1).

**Table 1:** Model metrics for covariate **albumin**

| Model | Train MSE | Test MSE | % of Patient Train $R^2$s for which Model > GPR | % of Patient Test $R^2$s for which Model > GPR | % of Patient 95% CIs for which Model is better than GPR | % of Patient 95% CIs for which Model is same as GPR |
| --- | --- | --- | --- | --- | --- | --- |
| GPR | 0.04 | 0.21 | — | — | — | — |
| HGP | 0.03 | 0.23 | 73.17 | 58.54 | 21.95 | 60.98 |
| MOE | 0.02 | 0.21 | 60.98 | 53.66 | 21.95 | 68.29 |

**Fig. 4:**
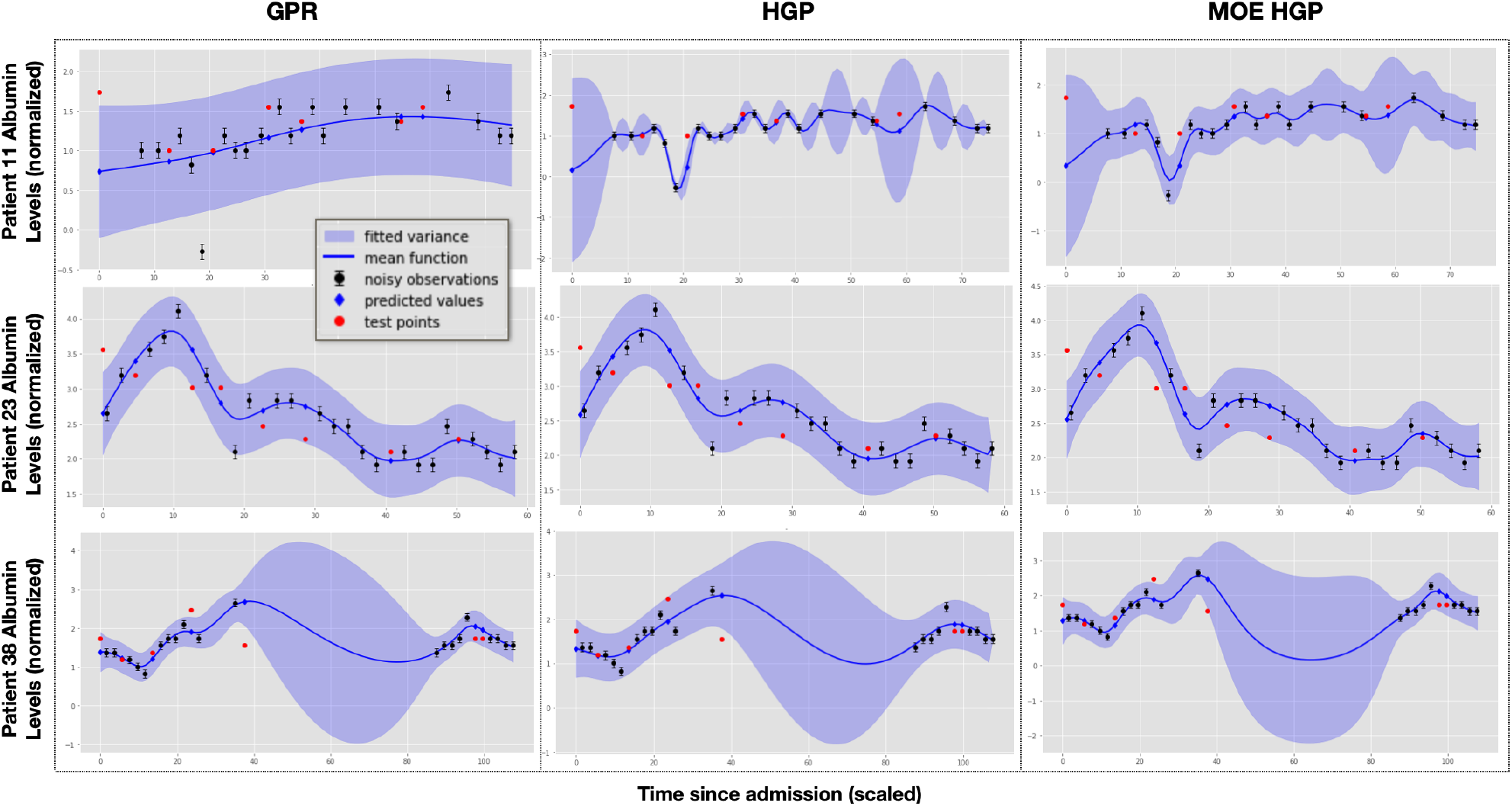
Cluster representative fits for covariate **albumin**. Patient 11 is a white male; Patient 23 is a Black female. Patient 38 is a white female.

We find substantial overlap in the the patient trajectories that benefit from the MOE HGP and HGP over GPR. Patient 7’s trajectory is a canonical case in which the *R*^2^ value is greatly improved with the HGP and MOE HGP (Fig. 5, Top). The mean function for GPR appears to a running average in the first half of the observed time points. The HGP and MOE HGP both provide better fits where GPR cannot. Similar to patient 7, patient 2’s trajectory has higher variance with GPR (Fig. 5, Bottom). This large variance has negative consequences on the 95% CI calibration. This patient has eight test points, so GPR gives a 95% CI of 0%, but the HGP and MOE HGP give 95% CIs of 25% since they each have two “outlier” test points. Taken together, these empirical results suggest that the two hierarchical models are more effective on these complex patient trajectories.

**Fig. 5:**
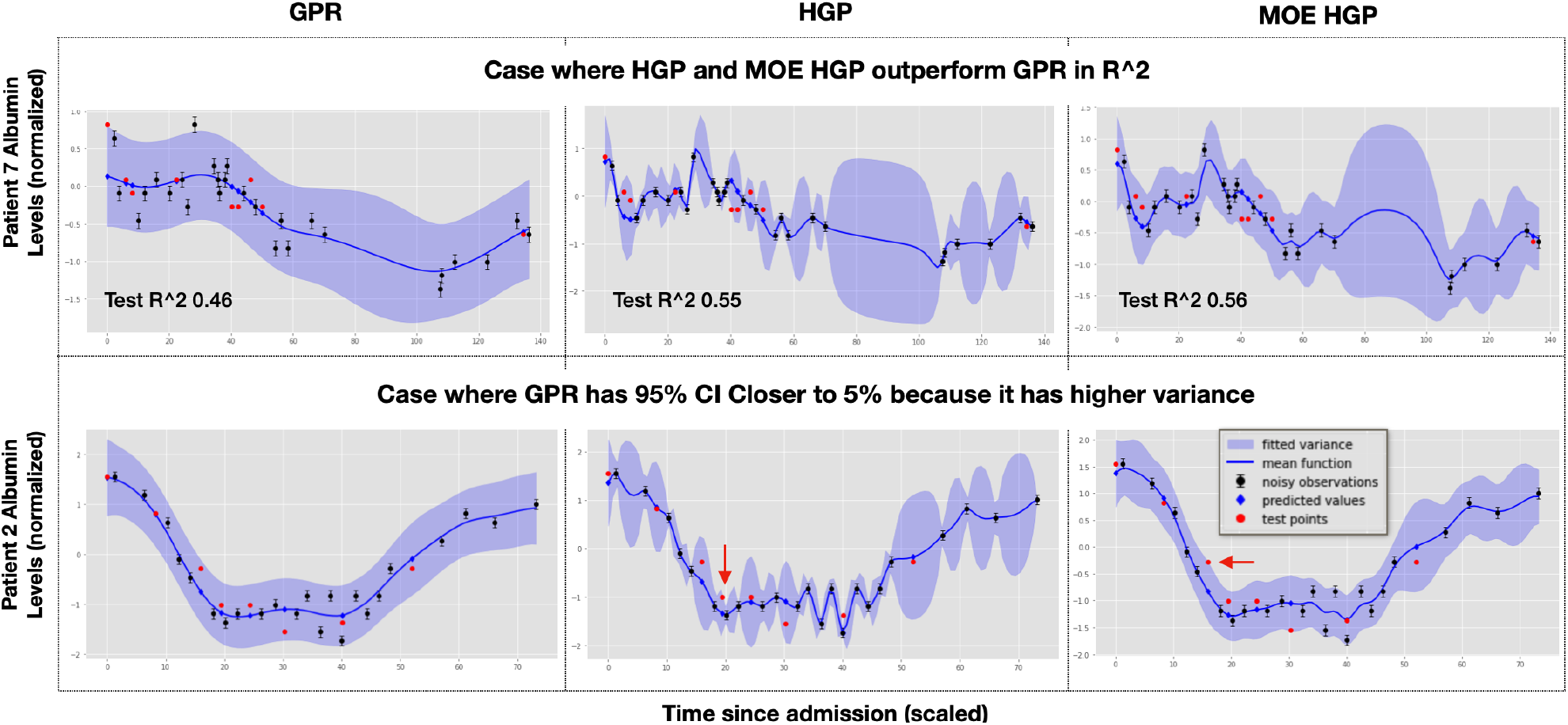
Exemplars of patient trajectories benefiting from HGP and MOE HGP.

The importance of group structure becomes more evident when we examine the kernel parameters at the patient level. The MOE HGP has lower spatial variance across all patients, as reflected in the distribution of the patient-level kernel variance parameters. GPR, lacking a group structure, defaults to learning a higher variance parameter. The structure of the MOE HGP is also useful for comparison across groups. When partitioning the patient cohort by ethnicity and sex, we see that Black patients have higher variance parameters than white patients do (Fig. 6). We do not observe meaningful differences in these parameters between male and female patients.

**Fig. 6:**
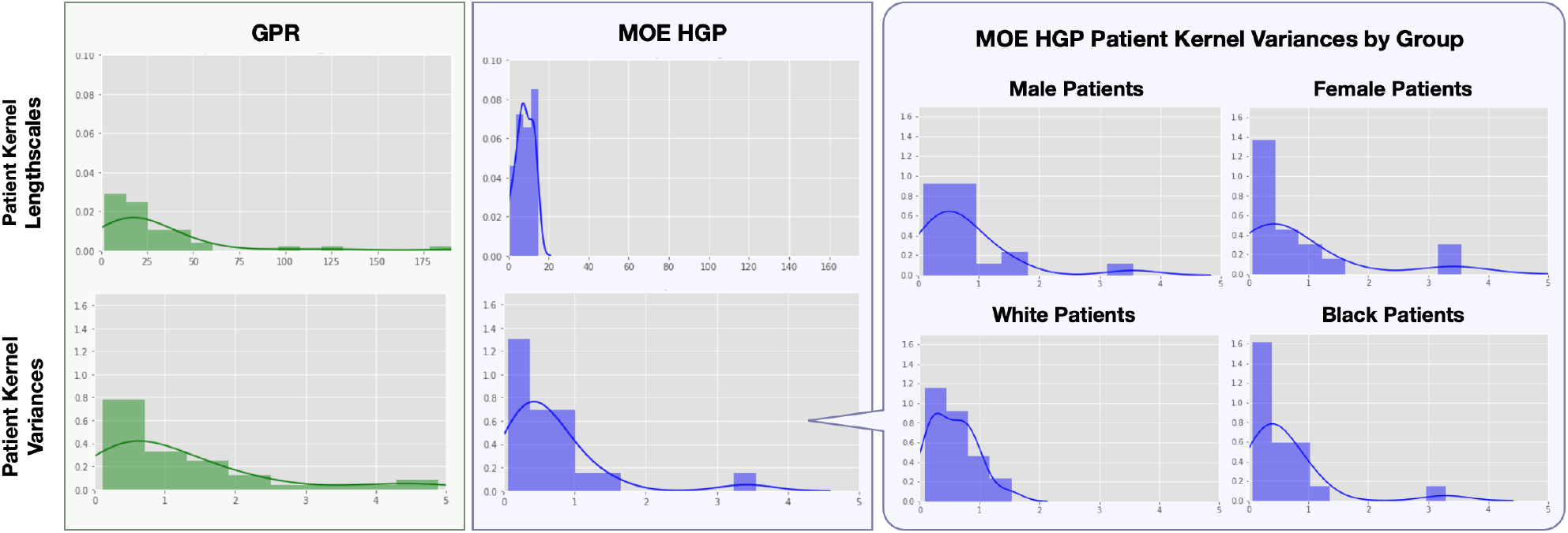
Patient level kernel parameters for GPR versus the MOE HGP for *albumin*.

Next, we fit the three models to the following clinical markers of COVID-19 disease progression for a randomly selected patient: *anion gap, creatinine, partial pressure of oxygen* (*PO*_2_ Arterial), *blood carbon dioxide levels* (*CO*_2_), *fraction of inspired oxygen* (*FIO*_2_) and *blood oxygen saturation* (Arterial *O*_2_ Content) (Fig. 7). Our experiments suggest that the MOE HGP effectively fits these markers for any randomly selected patient in the cohort.

**Fig. 7:**
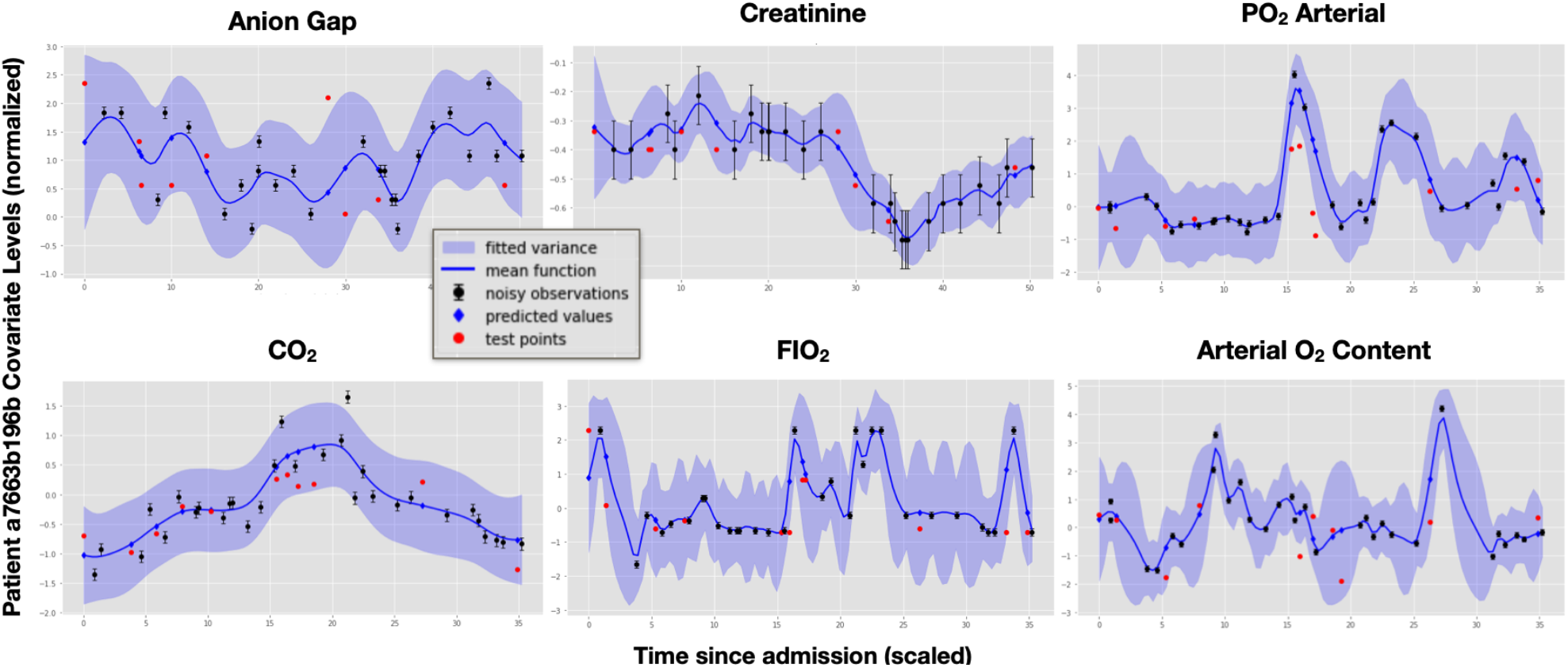
Covariate trajectories for a randomly selected patient in the cohort to demonstrate the robustness of the MOE HGP.

Across patients and groups, we see that the HGP and MOE HGP consistently outperform GPR in fitting patient trajectories for albumin, blood *CO*_2_, fraction of oxygen inspired *FIO*_2_, and lactic acid (Supplementary material Fig. 1-3, 5). These covariates – albumin as an indicator of kidney function and the remaining covariates as indicators of cardiovascular function – can inform immediate treatment decisions. Furthermore, the MOE HGP demonstrates superior uncertainty quantification over the HGP by giving the best 95% CI calibration at no observed cost to the test MSE, as reported for albumin, blood *CO*_2_, fraction of oxygen inspired *FIO*_2_ (Table 1, Supplementary material Tables 2-4). The MOE HGP’s strong performance, particularly in capturing complex trajectories with low spatial variance, can be attributed to its incorporation of group structures.

## 5. Conclusion

We propose a hierarchical mixture of experts Gaussian process (MOE HGP) model to fit and predict COVID-19 patient trajectories for clinically relevant covariates. We show that our MOE HGP model is effective in analyzing covariates and provides an in-depth analysis for albumin. We demonstrate the robustness of our model for an individual patient on indicators of blood oxygen levels like arterial *PO*_2_, *CO*_2_ and *FIO*_2_. Theses covariates are noisy yet useful for monitoring patient state in ICUs. Overall, the MOE HGP allows us to model groups separately while sharing signal across groups to enable more precise modeling of the natural group structure in patient populations without losing statistical power.

A natural extension of this work is to generalize the model to perform multi-output predictions. Because clinical covariates are often correlated, a multi-output GP that captures correlations between disparate covariates, in addition to correlations between observations within a single covariate, would be useful for more accurately modeling of clinical markers across time. With a multi-output model, we may include larger patient cohorts that are more diverse with respect to group attributes that could serve as proxies of socioeconomic status such as zip code, marriage status, and insurance status. We anticipate that we would be able to leverage such group structure to explore differences in disease trajectory or biases in treatment. Other group attributes like age, body mass index (BMI), and estimated glomerular filtration rate (eGFR) inform our understanding of how comorbidities such as obesity and renal disease impact disease progression within a certain socioeconomic or ethnic subpopulation.

Another direction of future work may be to apply contrastive learning, or methods that capture differences between the groups using parameters present in one but not the other.^24^ Contrastive modeling has been applied to linear dimension reduction^25^ and formalized to a probabilistic model-based alternative.^26,27^ With an extension of probabilistic contrastive modeling to Gaussian processes, we could improve the group-based prior for our model with information regarding differences between patients from traditionally marginalized populations, the “foreground” group, and their majority counterparts, the “background” group.

## 6. Acknowledgments and Appendices

We would like to thank the University of Pennsylvania Medical Center for providing the data and consultation regarding clinical domain knowledge. This work was funded in part by a COVID-19 grant from the Fast Grants program, a grant from the Helmsley Trust, a grant from the NIH Human Tumor Atlas Research Program, NIH NHLBI R01 HL133218, and NSF CAREER AWD1005627.

## Supplementary Material

### 1. Fitting Additional Covariates

We show the robustness of our mixture of experts hierarchical Gaussian process model by fitting and predicting patient trajectories on the following additional covariates: blood *CO*_2_, fraction of oxygen inspired *FIO*_2_, chloride, lactic acid, and creatinine. We present the results for albumin referenced in the main text as a point of reference. Again, we benchmark our mixture of experts (MOE) HGP model against Gaussian process regression (GPR) and a hierarchical Gaussian process (HGP).

#### 1.1. Blood CO_2_

**Fig. 1:**
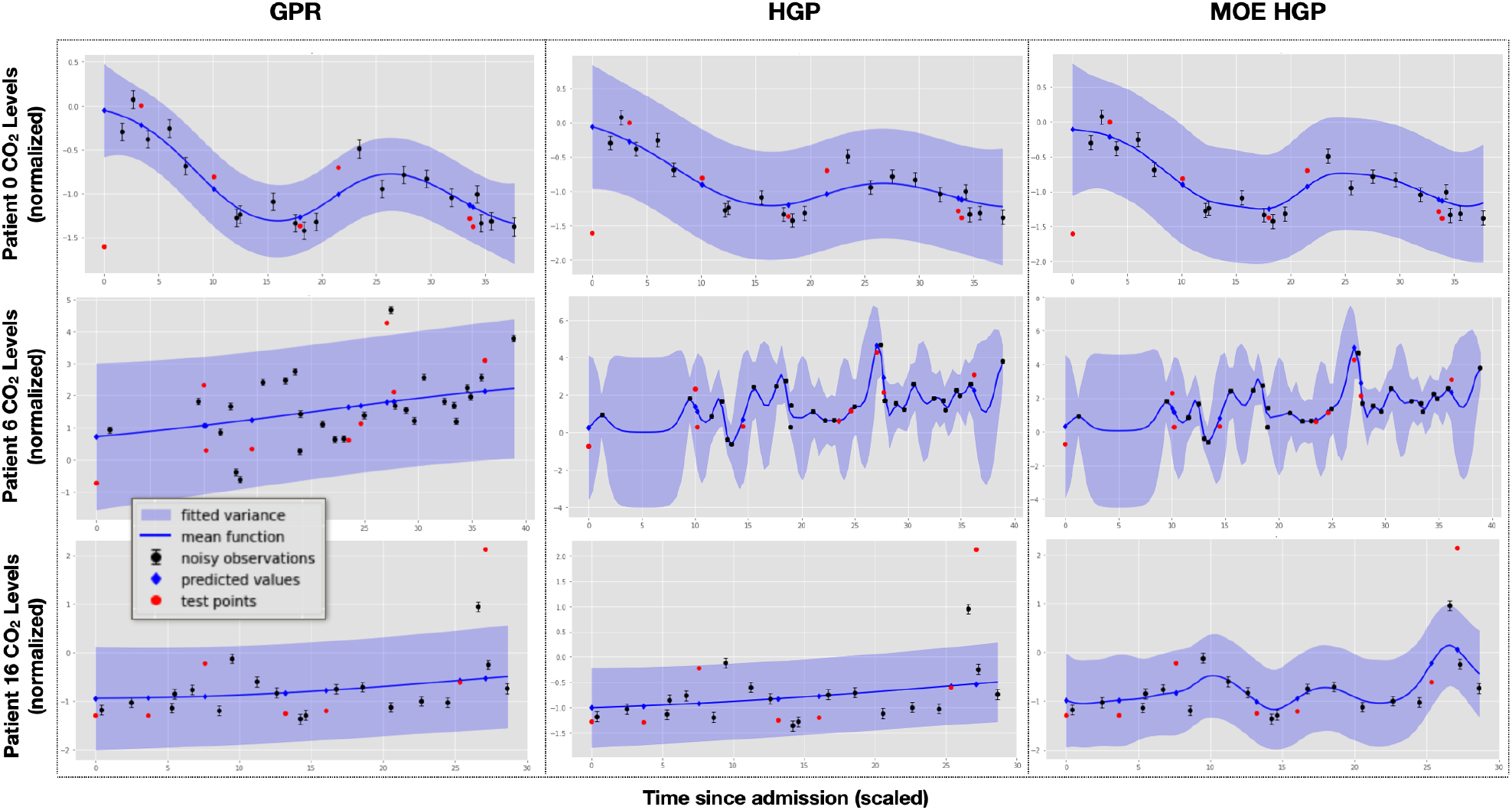
Cluster representative fits for *CO*_2_. Patient 0 is a representative of attributes 3 (ethnically Black). Patient 6 representative of attributes 0 (male) and 2 (ethnically white). Patient 16 is a representative of attribute 1 (female). The train/test MSEs for Patient 0 are 0.025*/*0.380 (GPR), 0.688*/*0.389 (HGP) and 0.024*/*0.352 (MOE). The train/test MSEs for Patient 6 are 1.00*/*1.51 (GPR), 0.126*/*0.463 (HGP) and 0.043*/*0.514 (MOE). The train/test MSEs for Patient 16 are 0.226*/*1.16 (GPR), 0.228*/*1.15 (HGP) and 0.081*/*0.729 (MOE).

**Table 1:** Model metrics for covariate *CO*_2_

| Model | Train MSE | Test MSE | % of Patient Train $R^2$s for which Model > GPR | % of Patient Test $R^2$s for which Model > GPR | % of Patient 95% CIs for which Model is better than GPR | % of Patient 95% CIs for which Model is same as GPR |
| --- | --- | --- | --- | --- | --- | --- |
| GPR | 0.20 | 0.70 | — | — | — | — |
| HGP | 0.09 | 0.61 | 47 | 47 | 24 | 65 |
| MOE | 0.058 | 0.57 | 76 | 47 | 29 | 65 |

#### 1.2. Fraction of Oxygen Inspired (FIO_2_)

**Fig. 2:**
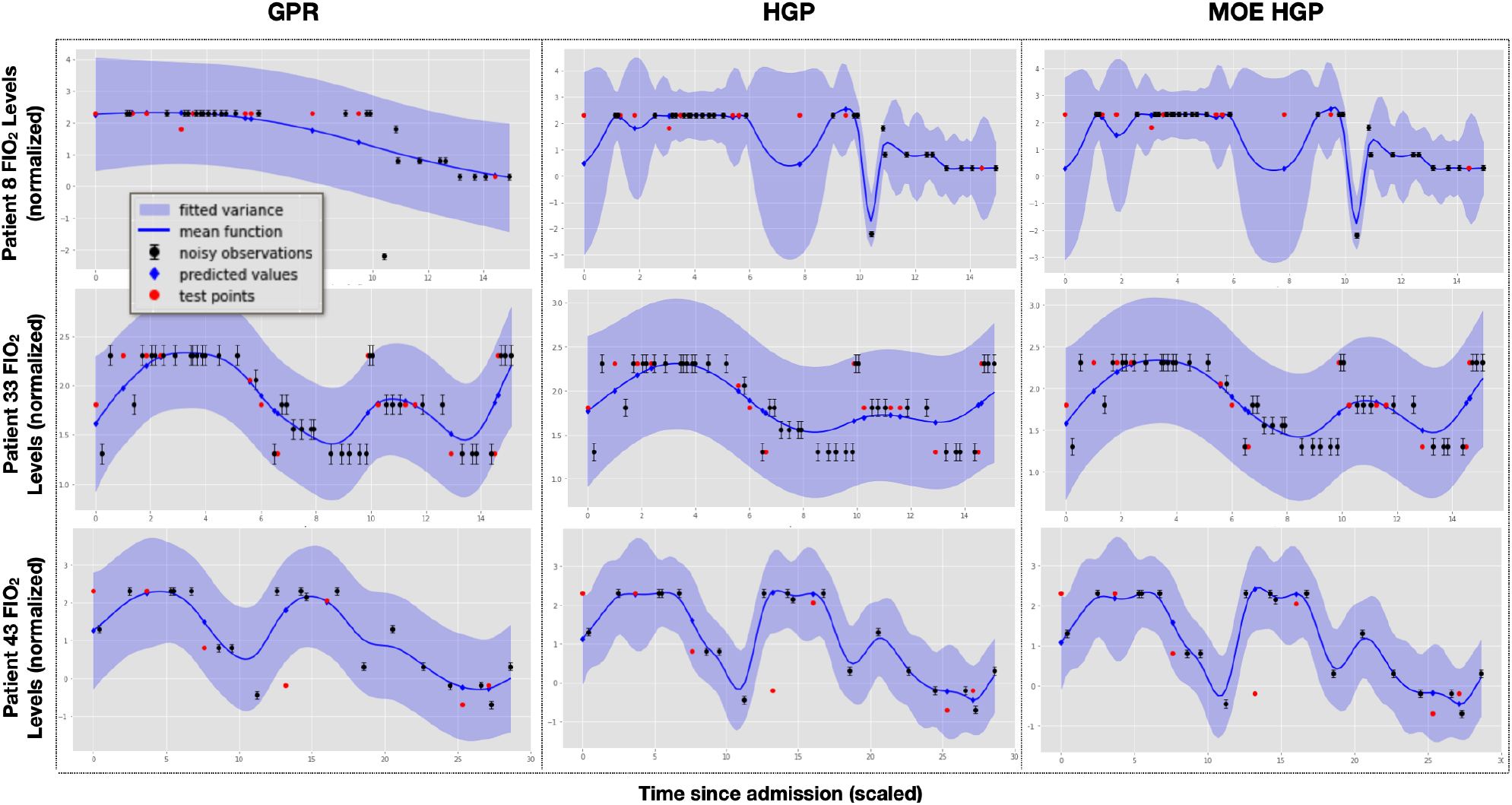
Cluster representative fits for *FIO*_2_. Patient 8 is a representative of attribute 0 (male). Patient 33 is a representative of attributes 1 and 3 (ethnically Black). Patient 43 is a representative of attribute 2 (ethnically white). The train/test MSEs for Patient 8 are 0.547*/*0.144 (GPR), 0.106*/*0.731 (HGP) and 0.043*/*0.907 (MOE). The train/test MSEs for Patient 33 are 0.053*/*0.090 (GPR), 0.323*/*0.105 (HGP) and 0.058*/*0.004 (MOE). The train/test MSEs for Patient 43 are 0.181*/*0.830 (GPR), 0.593*/*0.125 (HGP) and 0.026*/*0.133 (MOE).

**Table 2:** Model metrics for covariate *FIO*_2_

| Model | Train MSE | Test MSE | % of Patient Train $R^2$s for which Model > GPR | % of Patient Test $R^2$s for which Model > GPR | % of Patient 95% CIs for which Model is better than GPR | % of Patient 95% CIs for which Model is same as GPR |
| --- | --- | --- | --- | --- | --- | --- |
| GPR | 0.14 | 0.48 | — | — | — | — |
| HGP | 0.07 | 0.51 | 47 | 53 | 34 | 40 |
| MOE | 0.06 | 0.49 | 49 | 55 | 36 | 43 |

#### 1.3. Chloride

**Fig. 3:**
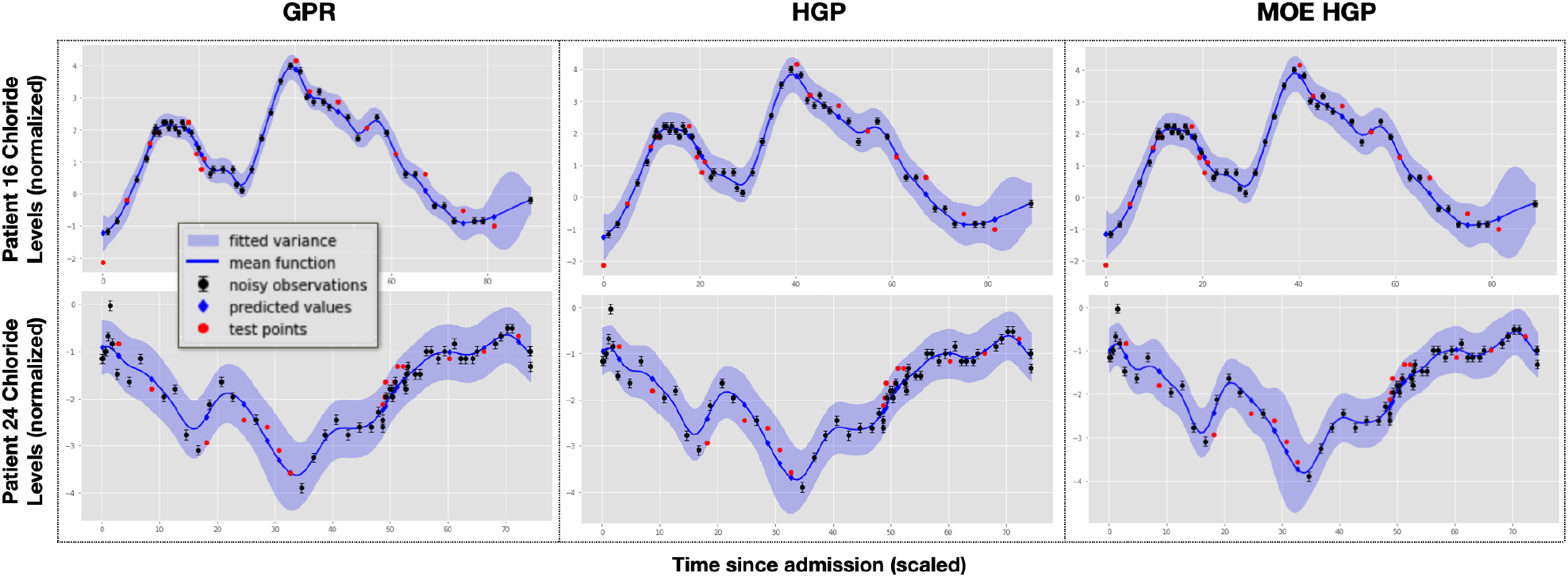
Cluster representative fits for **Chloride**. Patient 16 is a representative of attributes 1 (female) and 3 (ethnically Black). Patient 24 is a representative of attributes 0 (male) and 2 (ethnically white). The train/test MSEs for Patient 16 are 0.010*/*0.116 (GPR), 0.090*/*0.123 (HGP) and 0.016*/*0.128 (MOE). The train/test MSEs for Patient 24 are 0.049*/*0.074 (GPR), 0.024*/*0.077 (HGP) and 0.033*/*0.072 (MOE).

**Table 3:**
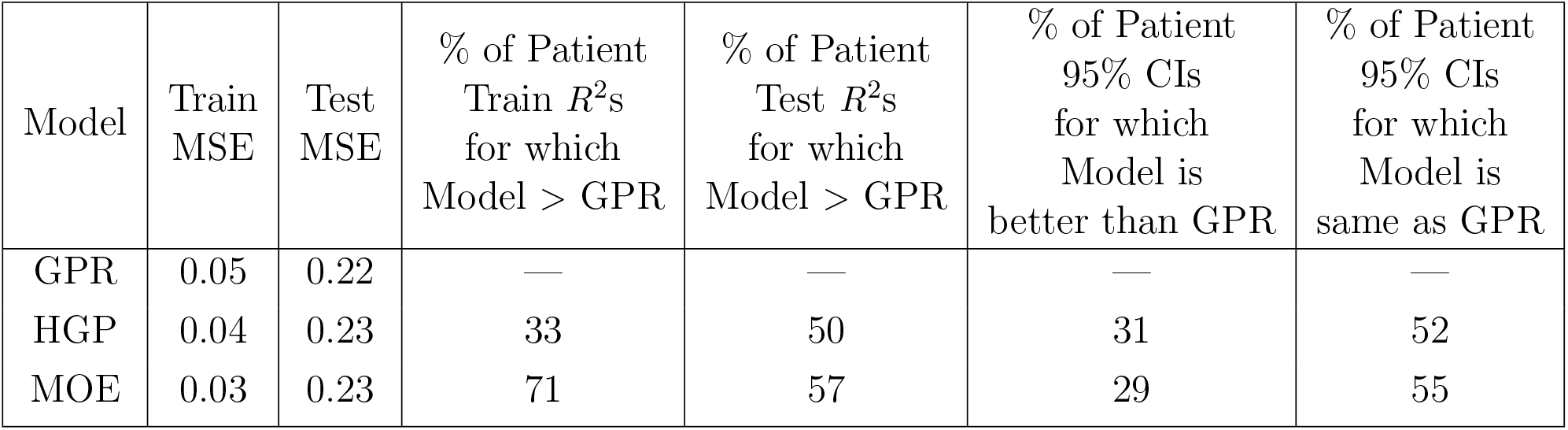
Model metrics for covariate **Chloride**

| Model | Train MSE | Test MSE | % of Patient Train $R^2$s for which Model > GPR | % of Patient Test $R^2$s for which Model > GPR | % of Patient 95% CIs for which Model is better than GPR | % of Patient 95% CIs for which Model is same as GPR |
| --- | --- | --- | --- | --- | --- | --- |
| GPR | 0.05 | 0.22 | — | — | — | — |
| HGP | 0.04 | 0.23 | 33 | 50 | 31 | 52 |
| MOE | 0.03 | 0.23 | 71 | 57 | 29 | 55 |

#### 1.4. Lactic Acid

**Fig. 4:**
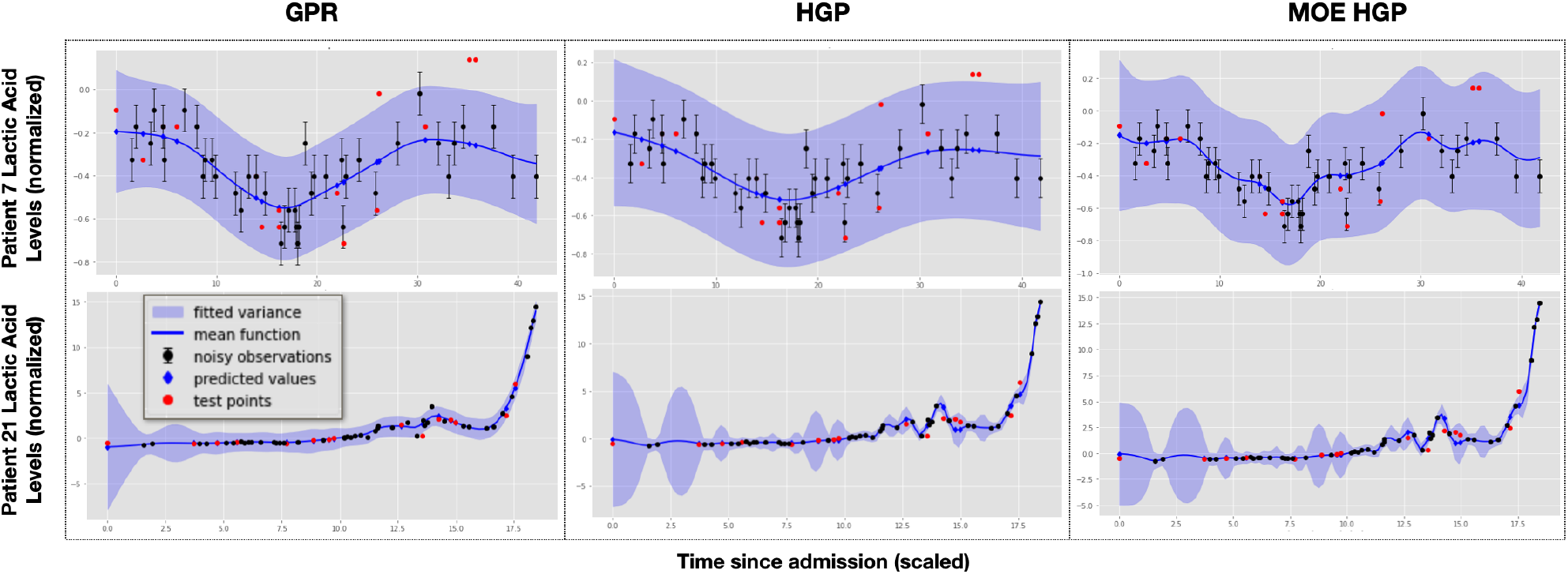
Cluster representative fits for **Lactic Acid**. Patient 13 is a representative of attributes 0 (male) and 2 (ethnically white). Patient 21 is a representative of attributes 1 (female) and 3 (ethnically Black). The train/test MSEs for Patient 13 are 0.012*/*0.040 (GPR), 0.055*/*0.042 (HGP) and 0.009*/*0.035 (MOE). The train/test MSEs for Patient 21 are 0.115*/*0.194 (GPR), 0.092*/*0.052 (HGP) and 0.180*/*0.052 (MOE).

**Table 4:** Fit on 33 COVID patients for **Lactic Acid**

| Model | Train MSE | Test MSE | % of Patient Train $R^2$s for which Model > GPR | % of Patient Test $R^2$s for which Model > GPR | % of Patient 95% CIs for which Model is better than GPR | % of Patient 95% CIs for which Model is same as GPR |
| --- | --- | --- | --- | --- | --- | --- |
| GPR | 0.06 | 0.19 | — | — | — | — |
| HGP | 0.02 | 0.22 | 47 | 59 | 21 | 59 |
| MOE | 0.01 | 0.22 | 55 | 53 | 24 | 56 |

#### 1.5. Creatinine

**Fig. 5:**
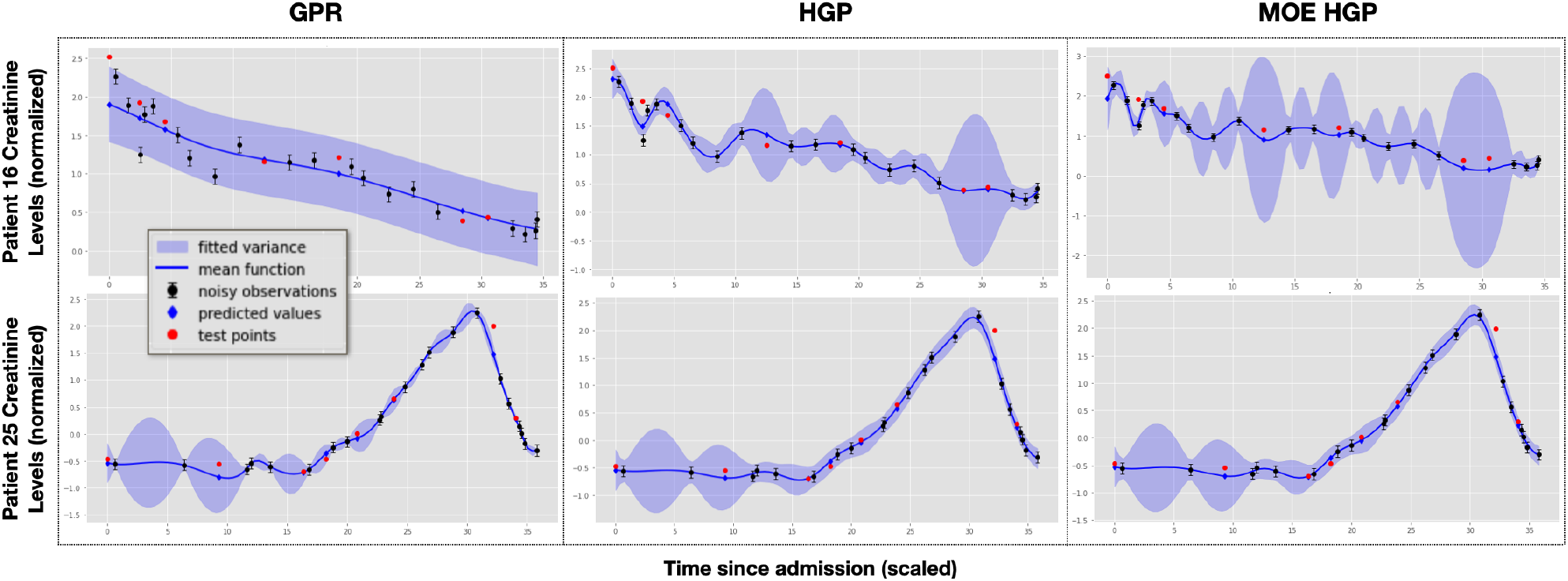
Cluster representative fits for **Creatinine**. Patient 1 is a representative of attribute 0 (male). Patient 16 is a representative of attribute 1 (female). Patient 25 is a representative of attribute 2 (ethnically white). Patient 36 is a representative of attribute 3 (ethnically Black). The train/test MSEs for Patient 16 are 0.037*/*0.070 (GPR), 0.104*/*0.043 (HGP) and 5.20 × 10^−4^*/*0.141 (MOE). The train/test MSEs for Patient 25 are 1.00 × 10^−5^*/*0.045 (GPR), 0.023*/*0.048 (HGP) and 8.10 × 10^−4^*/*0.039 (MOE).

**Table 5:** Model metrics for covariate **Creatinine**

| Model | Train MSE | Test MSE | % of Patient Train $R^2$s for which Model > GPR | % of Patient Test $R^2$s for which Model > GPR | % of Patient 95% CIs for which Model is better than GPR | % of Patient 95% CIs for which Model is same as GPR |
| --- | --- | --- | --- | --- | --- | --- |
| GPR | 0.05 | 0.07 | — | — | — | — |
| HGP | 0.003 | 0.13 | 51 | 47 | 27 | 53 |
| MOE | 0.001 | 0.14 | 89 | 44 | 29 | 44 |

## Footnotes

* *The code and supplementary material are available at: https://github.com/bee-hive/HGP-MOE

